# Diversity of sea star-associated densoviruses and transcribed endogenized viral elements of densovirus origin

**DOI:** 10.1101/2020.08.05.239004

**Authors:** Elliot W. Jackson, Roland C. Wilhelm, Mitchell R. Johnson, Holly L. Lutz, Isabelle Danforth, Joseph K. Gaydos, Michael W. Hart, Ian Hewson

**Affiliations:** Department of Microbiology, Cornell University, Ithaca NY USA; School of Integrative Plant Sciences, Bradfield Hall, Cornell University, Ithaca, NY USA; Department of Pediatrics, School of Medicine, University of California San Diego, La Jolla, California, USA; Scripps Institute, University of California San Diego, La Jolla, California, USA; SeaDoc Society, UC Davis Karen C. Drayer Wildlife Health Center – Orcas Island Office, Eastsound, WA 98245 USA; Department of Biological Sciences, Simon Fraser University, Burnaby, British Columbia V5A 1S6, Canada

## Abstract

A viral etiology of Sea Star Wasting Syndrome (SSWS) has been largely explored using metagenomics leading to the conclusion that a densovirus is the predominant DNA virus associated with this syndrome, and, thus, the most promising viral candidate pathogen. Single-stranded DNA viruses are however highly diverse and pervasive among eukaryotic organisms which we hypothesize may confound the association between densoviruses and SSWS in sea stars. To test this hypothesis and assess the association of densoviruses to SSWS, we compiled past metagenomic data with new metagenomic-derived viral genomes from sea stars collected from Antarctica, California, Washington, and Alaska. We used 179 publicly available sea star transcriptomes to complement our approaches for densovirus discovery. Lastly, we focus the study to SSaDV, the first sea star densovirus discovered, by documenting its biogeography and putative tissue tropism. Transcriptomes contained mostly endogenized densovirus elements similar to the NS1 gene, while >30 complete and near-complete densoviral genomes were recovered from viral metagenomes. SSaDV was associated with nearly all tested species from southern California to Alaska, and in contrast to previous work, we show SSaDV is one genotype among a high diversity of densoviruses present in sea stars across the west coast of the United States and globally that are commonly associated with grossly normal (i.e. healthy or asymptomatic) animals. The diversity and ubiquity of these viruses in wild sea stars confounds the original hypothesis that one densovirus was the etiologic agent of SSWD.

**Importance:** The primary interest in sea star densoviruses, specifically SSaDV, has been their association with Sea Star Wasting Syndrome (SSWS), a disease that has decimated sea star populations across the west coast of the United States since 2013. The association of SSaDV to SSWS was originally drawn from metagenomic analyses concluding that it was (1) the only densovirus present in the metagenomic data and (2) the most likely viral candidate based on representation in symptomatic sea stars. We reassessed the original metagenomic data with additional genomic datasets and found that SSaDV was one of ten densoviruses present in the original dataset and was no more represented in symptomatic sea stars than in asymptomatic sea stars. Instead, SSaDV appears to be a widespread, generalist virus that exists among a large diversity of densoviruses present in sea star populations.

## Introduction

Single-stranded (ss) DNA viruses are among the most diverse and prevalent group of viruses infecting eukaryotes, bacteria, and archaea (1–4). Recognition of their ubiquity has been made possible through the use of rolling circle amplification that preferentially amplifies circular nucleic acid templates prior to high-throughput sequencing (5, 6). As a result, ssDNA viruses that possess circular genomes are significantly overrepresented compared to those with linear genomes. There are currently nine established families of ssDNA viruses that infect eukaryotes, only two of which possess linear genomes - *Bidnaviridae* and *Parvoviridae* (7). While our knowledge of the circular ssDNA viruses has expanded tremendously, the discovery of linear ssDNA viruses has lagged. Collectively, the known viral diversity of these two families likely represents only a small proportion of actual extant diversity, particularly within the subfamily *Densovirinae* (family *Parvoviridae*).

Currently 17 viral species are recognized by the International Committee on the Taxonomy of Viruses that belong to the subfamily *Densovirinae* (commonly referred to as densoviruses) (8). Densoviruses infect invertebrates and until recently, were only known to infect insects (orders Blattodea, Diptera, Hemiptera, Hymenoptera, Lepidotera, and Orthoptera) and decapod crustaceans (shrimp and crayfish) (9). Unlike circular ssDNA virus groups, the discovery of novel linear densoviruses genomes has occurred primarily via the use of classical methods, such as viral purification through cell culture or viral enrichment via an animal model followed by nucleic acid sequencing, but these classical methods are more recently supplemented by the use of high-throughput sequencing to explore viral diversity. Transcriptomic and metagenomic analyses have generated a growing body of evidence to suggest that densoviruses infect a more phylogenetically diverse array of invertebrate hosts outside of the phylum Arthropoda (10–13). Expanding the known potential host range of these viruses will lead to insights into biology and evolution of densoviruses and the hosts they infect.

The 2014 discovery of densoviruses associated with sea stars and sea urchins (phylum: Echinodermata) was a significant step towards expanding the range of densoviruses beyond arthropods (11, 13, 14). The primary interest in echinoderm densoviruses has been their association with Sea Star Wasting Syndrome (SSWS; also referred to as Sea Star Wasting Disease and Asteroid Idiopathic Wasting Syndrome) which is an epidemic that affected sea stars on the east and west coast of North America (11, 15). Currently, the etiology of SSWS is unknown. A densovirus, commonly known as <u>s</u>ea <u>s</u>tar <u>a</u>ssociated <u>d</u>ensovirus or SSaDV, was hypothesized to cause SSWS but it has yet to be determined if SSaDV is a pathogen (11, 16). Originally, SSaDV was thought to be the etiological agent of SSWS observed in both Pacific and Atlantic sea star populations (11, 15). However, the recent discovery of a genetically similar second densovirus associated with sea stars on the Atlantic coast, *<u>A</u>sterias <u>f</u>orbesi* <u>a</u>ssociated <u>d</u>ensovirus or AfaDV, raises new questions about densovirus diversity and the putative role of densoviruses in SSWS (13). The discovery of AfaDV prompted us to explore the diversity of densoviruses at a greater geographic scale and to define the biogeography of SSaDV and expand our understanding of the ecology, evolution, and diversity of these viruses.

Here, we employ a multi-omic approach to document the biodiversity of densovirus populations in sea stars by first reassessing the original metagenomic dataset leading to the association of densoviruses and SSWS. We used publicly available sea star transcriptomes/genomes and sea star viral metagenomes for viral discovery, and PCR to document the prevalence, putative tissue tropism, and biogeography of SSaDV. We report the discovery of >30 novel sea star densoviruses associated with sea stars from the Southern Ocean around Antarctica (11 genomes) and from the temperate eastern Pacific (24 genomes), and the observation of numerous endogenized viral elements (EVEs) from sea star transcriptomes and genomes. We found that SSaDV putatively has a wide tissue tropism and is associated with sea stars in the eastern Pacific across a broad latitudinal range from southern California to Alaska, corroborating previous findings by Hewson *et al*., 2014 (11). The identification of SSaDV as one among many densoviruses infecting sea stars on the west coast of the United States, suggests that densoviruses may comprise a normal component of the sea star microbiome, bringing into question the association of densoviruses to SSWS.

## Methods

### Tissue Collection & DNA Extractions

We collected 991 tissue samples from 661 individual sea stars spanning 11 species from 41 locations from the temperate eastern Pacific coast of the United States from 2005, 2014-2019 (**Supplemental Table 1**). The majority (96.5%, n=957) of tissue samples were collected from 2013-2019. Thirty samples were collected during the peak of SSWS epidemic observed from mid 2013-2015 in the Northeast Pacific (17). The majority of samples collected were also from asymptomatic individuals (85% of sea stars sampled). Tissues were collected from sea star specimens by either vivisection immediately upon collection, then flash frozen in liquid nitrogen upon collection and stored at −80°C or −20°C until dissection, or sampled from individuals in the field non-lethally and preserved in RNAlater (SigmaAldrich) or EtOH (**Supplemental Table 1**). Coelomic fluid samples were collected only from vivisected specimens using a 25G x 1. (0.5mm x 25mm) needle attached to a 3 mL syringe inserted through the body wall into the coelomic cavity. DNA was extracted from tissues and coelomic fluid using the Zymo Research Quick-DNA Miniprep Plus kit or the Zymo Research Duet DNA/RNA Miniprep kit following the manufacture’s protocol. DNA was quantified using a Nanodrop or Quant-iT PicoGreen dsDNA Assay kit (Invitrogen) (**Supplemental Table 1**).

### PCR and Sanger Sequencing of SSaDV

Primers were designed targeting the structural gene of SSaDV using Primer3web (version 4.1.0). The primers were as follows: VP1 forward primer (5’-TGGCCACTCATCATGTCTCT-3’) and VP1 reverse primer (5’ – CTTGGGGTCCTTCATGAGC – 3’). NEB Q5 High-Fidelity DNA Polymerase was used following the manufacturer’s protocol for a 50μl reaction volume. Thermal cycling was performed in a Bio-Rad C1000™ Thermal Cycler using the following conditions: initial denaturing at 98°C for 30 seconds followed by 35 cycles of denaturing (98°C for 10 seconds), annealing (67°C for 20 seconds), and extension (72°C for 20 seconds) followed by a final extension of 72°C for 2 minutes. Annealing temperature was based on NEB Tm Calculator recommendation using default primer concentration of 500nM. The resulting amplicon of the PCR reaction was 534 nucleotides (nt). All PCR reactions included a positive control, a kit negative control, and PCR reagent negative control to account for false positives and false negatives. 10-15μl of a PCR reaction was used for gel visualization. The remaining PCR product was processed using a ZR-96 DNA Clean & Concentrator-5 kit (Zymo Research) and submitted to Cornell Core Biotechnology Resource Center Genomics Facility for DNA sequencing. DNA sequencing was performed on Applied Biosystems Automated 3730xl DNA Analyzers using Big Dye Terminator chemistry and AmpliTaq-FS DNA Polymerase.

### Viral Metagenome Preparation

Five RNA viral metagenomes were prepared from five different species of sea stars - *Pisaster ochraceus* (n=1), *Labidiaster annulatus* (n=1), *Leptasterias* spp. (n=1), *Mediaster aequalis* (n=1), and *Neosmilaster georgiansus* (n=1) (**Supplemental Table 2**). The preparation of viral metagenomes followed Hewson *et al*., 2018 modified from Thurber *et al*., 2009 (16, 18). Pyloric caeca from sea stars were homogenized in a 10% bleached-cleaned NutriBullet with 0.02μm-filtered 1X PBS. Tissue homogenates were pelleted by centrifugation at 3,000 x g for 5 minutes, and the supernatant was syringe filtered through Millipore Sterivex-GP 0.22μm polyethersulfone filters into 10% bleach-treated and autoclaved Nalgene Oak Ridge High-Speed Centrifugation Tubes. Filtered homogenates were added to a 10% (wt/vol) PEG-8000 in 0.02μm-filtered 1X PBS with a final volume of 35mL and precipitated for 20 hours at 4°C. Precipitated nucleic acids, and other cell material, were pelleted by centrifugation at 15,000 x g for 30 minutes. The supernatant was decanted and pellets were resuspended in 2mL of 0.02μm-filtered 1X PBS. Half of the sample (1mL) was treated with 0.2 volumes (200μl) of CHCl3, inverted three times and incubated at room temperature for 10 minutes. After a brief centrifugation, 800μl of supernatant was transferred into 1.5mL microcentrifuge tube. Samples were treated with 1.5μl of TURBO DNase (2U/μl) (Invitrogen), 1μl of RNase One (10U/μl) (Thermo Scientific), and 1μl of Benzonase Nuclease (250U/μl) (MilliporeSigma) and incubated at 37 °C for 3 hours. 0.2 volumes (160μl) of 100mM EDTA was added to the sample after incubation. Viral RNA was extracted using the ZR Viral RNA kit (Zymo Research). RNA was converted into cDNA and amplified using the WTA2 kit (Sigma-Aldrich). Prior to sequencing, samples were processed using a ZR-DNA Clean & Concentrator-5 kit, and DNA was quantified by Quant-iT PicoGreen dsDNA Assay Kit. Samples were prepared for Illumina sequencing using the Nextera XT DNA library preparation kit prior to 2×250bp paired-end Illumina MiSeq sequencing at the Cornell Core Biotechnology Resource Center Genomics Facility.

### Metagenome-derived Viral Genome Discovery

Raw paired-end reads were quality trimmed to remove Illumina adapters and phiX contamination. Reads were merged and normalized to a target depth of 100 and a minimum depth of 1 with an error correction parameter. Read quality filtering, trimming, contamination removal, merging, normalization, and read mapping were performed using the BBtools suite (19). Both merged and unmerged reads were used for *de novo* assembly using SPAdes v 3.11.1 (20). Contigs shorter than 3000 nt were discarded after assembly and the remaining contigs were subjected to tBLASTx against a curated in-house database containing 453 genomes from all nine families of eukaryotic ssDNA viruses. Contigs with significant sequence similarity at *e*-value < 1×10^−8^ to a densovirus genome were reviewed in Geneious version 9.1.5 (21). Open reading frames (ORFs) were called in Geneious using a minimum size of 550 nt with a standard genetic code and a start codon of ATG. Hairpin structures were identified using Mfold (22). After verifying the contigs as densovirus sequences, reads were mapped back to contigs with a minimum identity of 0.95 to obtain average read coverage and total reads mapped to contigs (**Supplemental Table 3**). All densovirus sequences have been deposited in GenBank (**Table 1**). In addition to the 5 viral metagenome libraries sequenced in this study, we reanalyzed 30 DNA and 21 RNA viral metagenomes published elsewhere (NCBI BioProjects: PRJNA253121, PRJNA417963, PRJNA637333) using the assembly approach described above (**Supplemental Table 2**) (11, 16).

### Sea Star Transcriptome Analysis

A total of 179 sea star RNA-seq paired-end libraries were downloaded from NCBI (**Supplemental Table 4**). FastX was used to remove reads with lengths <50 nt and a quality score <30 (23). Trimmomatic was used to trim adapters (24). Libraries were assembled using default parameters in Trinity v2.1.1 (25). Assembled contigs were annotated against a protein database of sea star densoviruses using DIAMOND with an e-value cutoff of <1 x 10^−5^ (26). Contigs containing sequences with significant similarity to the sea star associated densovirus protein database were isolated, ORFs were called, and amino acid sequences from ORFs were further checked by BLASTp against the NCBI non-redundant database. Top BLAST results for each densovirus-like sequence against the NCBI non-redundant database were downloaded for amino acid MUSCLE alignment and visualized using Geneious (27, 21). To verify if the sequences were from the host, BLASTn was performed querying isolated densovirus-like sequences against available sea star genomes (*Asterias rubens*, NCBI taxid: 7604, and *Acanthaster planci*, 133434).

### Densovirus Phylogenetics

Phylogenetic analysis was performed on 89 densoviruses sequences that included 39 sea star-associated densoviruses and 50 densovirus sequences from complete or near-complete genomes available on NCBI. All densovirus sequences included in the phylogeny were from extant viruses (i.e., no endogenized densovirus elements). Amino acid sequences from the NS1 gene were aligned with MUSCLE using default parameters (27). The region of NS1 used for alignment (sequence length of 434.9 ± 40.2 (mean ± standard deviation)) spanned motif I of the replication initiation motifs past Walker C of the Walker box ATPase motifs. Phylogenetic relationships between densovirus genomes were inferred by a LG + G + I + F substitution model selected by smart model selection (SMS) in PhyML 3.0 (28). Branch support was determined by bootstrapping for 100 iterations. The resulting maximum likelihood phylogenetic tree was visualized and annotated using iTOL (29). CD-HIT was used to identify viral species using a 85% amino acid sequence identity of NS1 (30, 31).

## Results

### Reanalysis of metagenomes published in Hewson *et al*., 2014

The reanalysis of the viral metagenomic data presented in Hewson *et al*., 2014 (11), led to the discovery of 9 additional densovirus genomes in addition to SSaDV (**Figure 1**, **Table 1**). The densovirus contigs ranged in nucleotide length from 3391 to 6053 nt (5002 ± 921 (mean ± standard deviation)). SSaDV was the only densovirus assembled into a complete or near complete genome across multiple metagenomes and had the highest read recruitment among all libraries, which is likely why it was the only densovirus contig produced in the previously performed global assembly (11) (**Figure 1, Table 1, Supplemental Table 3**). The previously published partial SSaDV genome (5,050 nt) lacked the NS3 ORF and inverted terminal repeats (ITRs). In this study, we recovered three SSaDV genomes of varying sizes from three of the 32 metagenomes (**Table 1**). The largest of these genomes (6,053 nt) contained the expected ORFs (NS1, NS2, NS3, and VP), ITRs, and hairpins within the ITRs, and therefore likely represents a complete genome. It is possible that the ITR region of the genome is not complete, due to challenges posed by assembling regions with high frequency of repeats using short read technology. Members of the genus *Ambidensovirus* typically have ITRs >500 nt, which is considerably longer than the ITR regions we observed (9). The length of the ITRs in SSaDV were 260 nt on both sides of the genome and contained canonical hairpin structures that were 223 nt and are thermodynamically favorable (ΔG = −106.40).

**Figure 1:**
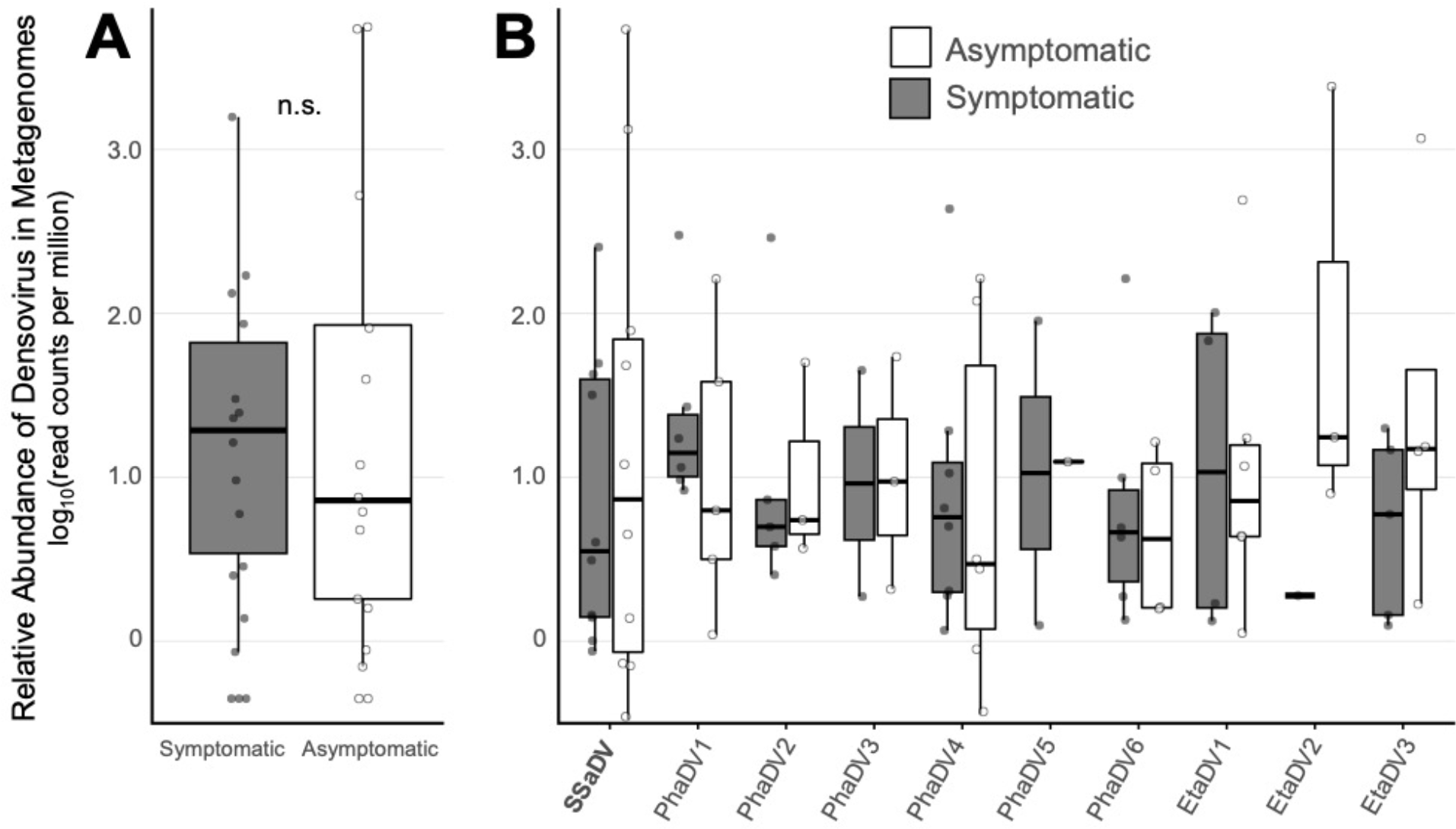
Reanalysis of metagenomic data presented in Hewson *et al*., 2014. SSaDV is one of ten densoviruses present in the data set and based on read mapping analysis (≥95% read identity) and is not more abundant in symptomatic compared to asymptomatic individuals. (A) Relative abundance of all reads recruited to densovirus genomes. N.S. (no significance) based on welch two sample t-test (*p* = 0.7697, df = 25.137, t = −0.29592). (B) Read recruitment separated by densovirus genotype.

### SSaDV biogeography and tissue tropism

A total of 146 of 661 animals were virus positive for SSaDV based on the PCR assay, equating to a global prevalence of 22.1% (**Figure 2, Supplemental Table 1**). No samples were PCR positive from tissues collected in 2005. A total of 123 of 146 PCR amplicons were successfully Sanger sequenced, all confirming the specificity of the PCR assay (**Supplemental Table 1**). Only 22 of 99 (22.2%) symptomatic (i.e. SSWS affected) sea stars were PCR positive, and 124 of 562 (22.0%) asymptomatic sea stars tested PCR positive. SSaDV was detected in 28 of the 41 locations that spanned a broad latitudinal range in the eastern Pacific from southern California to southeastern Alaska (**Figure 2**). Eight of 11 sea star species tested positive, which include the following species (virus positive / sample total): *Pisaster ochraceus* (88/287)*, Pisaster brevispinus* (11/11)*, Pisaster giganteus* (3/9), *Pycnopodia helianthoides (2/72), Evasterias troschelii* (27/109)*, Dermasterias imbricata* (8/34), *Henricia spp*. (2/17), *Leptasterias spp*. (4/33), and *Patiria miniata* (1/85). Only one animal was tested for each of the three species in which SSaDV was not detected. These included *Orthasterias koehleri*, *Pteraster tesselatus*, and *Solaster stimpsoni*.

**Figure 2:**
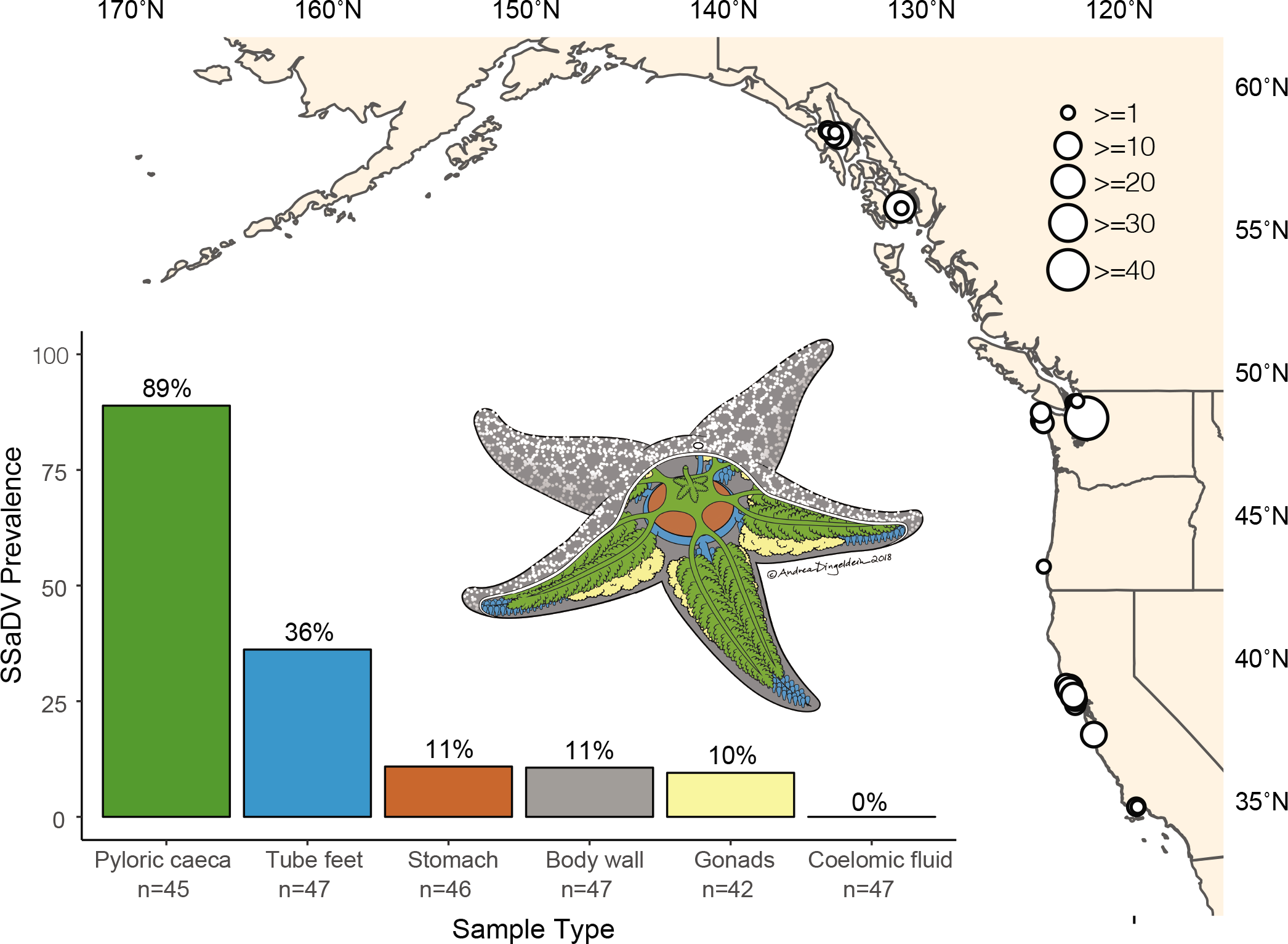
SSaDV is broadly distributed across the northeastern Pacific Ocean and putatively has a wide tissue tropism. White dots on map indicated PCR positive sample, and the size of the dot corresponds to total number of PCR positive samples at each site. Tissue tropism assessed from 3 sea star species collected from one site (Langley Harbor, Washington). Colors of each bar corresponds to the anatomical region in the sea star illustration. Prevalence defined as the number of PCR positive samples divided by the total number of samples tested for each tissue.

Fine dissections of *Pisaster ochraceus* (n= 26 individuals), *Evasterias troschelii* (n = 10), and *Pisaster brevispinus* (n = 11) collected from Langley Harbor, Washington were used assess putative tissue tropism. The viral prevalence among tissues was calculated by the number of tissues positive divided by the total number of tissues collected between these three species. SSaDV was detected most frequently in the pyloric caeca (89%, 40/45) followed by tube feet (36%, 17/47), stomach (11%, 5/46), body wall (11%, 5/47), and gonads (10%, 4/42) (**Figure 2**). Similar to AfaDV, SSaDV was not detected in the coelomic fluid (0%, 0/47) (13).

### Genome discovery, genome comparison, motif annotation, and phylogeny

An additional 29 densovirus genomes were recovered from newly prepared and reanalyzed sea star metagenomes (**Table 1**). The densovirus contigs ranged in size from 3061 to 5963 nt (5179.0 ± 684.4 (mean ± standard deviation Most densovirus-containing contigs (n = 28) corresponded to near complete genomes containing all the expected ORFs but lacking either ITRs or hairpins with the ITRs. The average size of the ORFs found in sea star densovirus were as follows (± standard deviation): NS1 1694.8 nt (± 33.3), NS2 883.1 nt (± 28.5), NS3 807.8 (± 109.2), and VP 2723.0 nt (± 98.2). The pairwise nucleotide identity was greater than amino acid (aa) identity among sea star densovirus genomes for NS1, NS3, and VP ORFs (**Figure 3**). The NS1 ORF had the highest sequence conservation (55.7% average nt and 43.2% average aa pairwise identity) compared to NS3 (32.6% average nt and 18.6% aa acid pairwise identity), and VP (43.8% average nt and 34.2% average aa pairwise identity). The current delineation for a new parvovirus species is based on the pairwise amino acid sequence identity of NS1. Parvoviruses encoding for NS1 proteins with a >85% pairwise amino acid sequence identity are considered the same viral species (31). Using this species definition, 29 new sea star densovirus species were defined from the 39 genotypes discovered. There were 8 viral species that contained 2 or 3 genotypes, and 21 species contained a single genotype. (**Table 1**).

**Figure 3:**
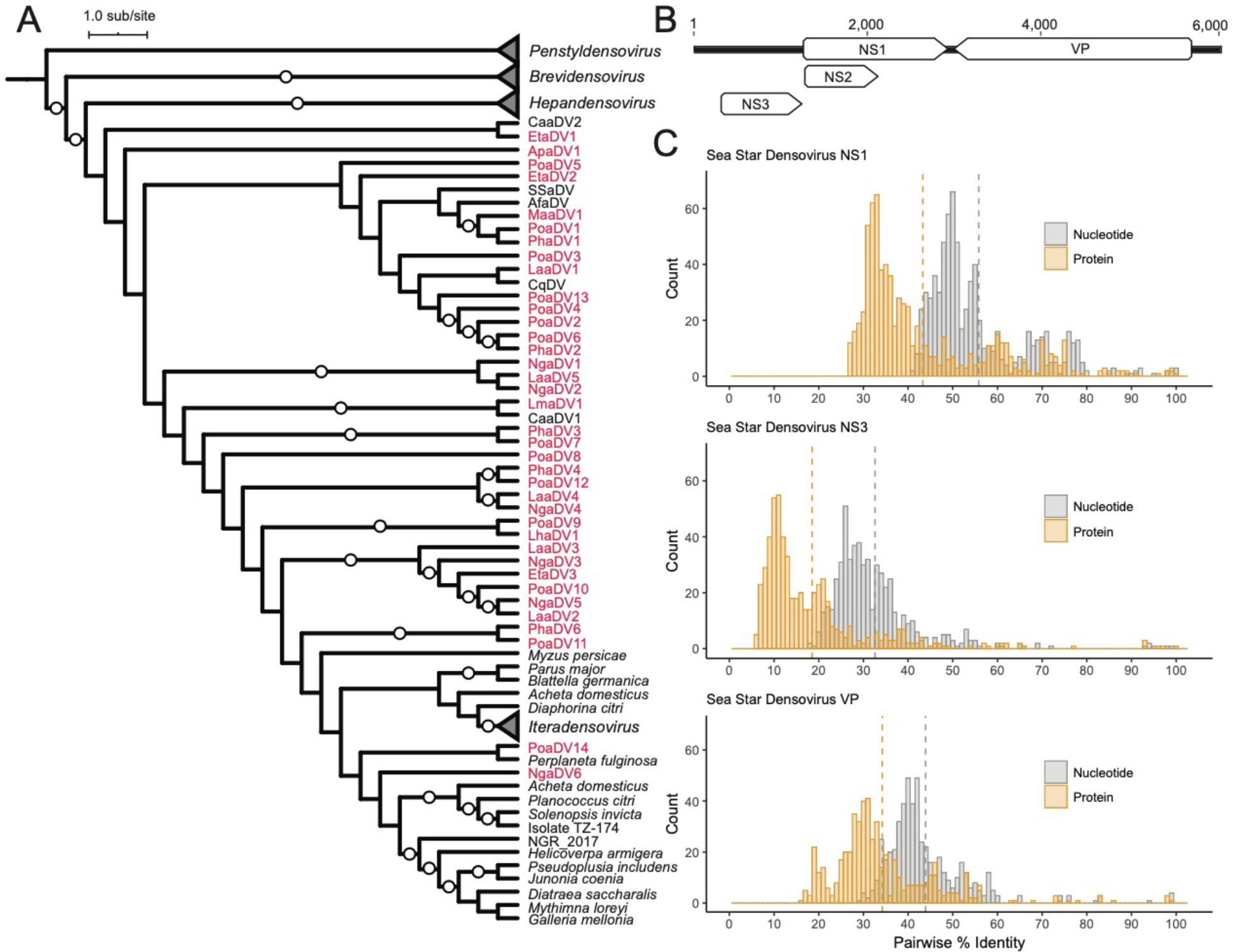
Sea star-associated densoviruses are genetically diverse and are not monophyletic. (A) Cladogram of a maximum likelihood phylogenetic tree of densoviruses based on alignment of amino acid sequences from NS1 gene. Collapsed nodes represent densovirus genera while all other branches belong to the genus *Ambidensovirus*. Red names indicate genomes discovered in this study. White circles represent 90-100% bootstrapped support. (B) Representative densovirus genome showing genome organization. (C) Histograms of nucleotide and amino acid pairwise identity comparisons between all sea star-associated densoviruses for NS1, NS3, and VP ORFs. Dotted lines indicate mean pairwise identity.

All sea star densoviruses discovered thus far have ambisense genomes that fall into subgroups A and B, which differ only by the VP ORF organization (32) (**Figure 4**). The NS1 and VP ORFs identified in this study contain all the expected motifs that are characteristic of densoviruses (32). These motifs include: RCR I and RCR II of the replication initiation motifs, Walker A, B, and C of the NTP-binding and helicase motifs, and the viral phospholipase A2 motif.

**Figure 4:**
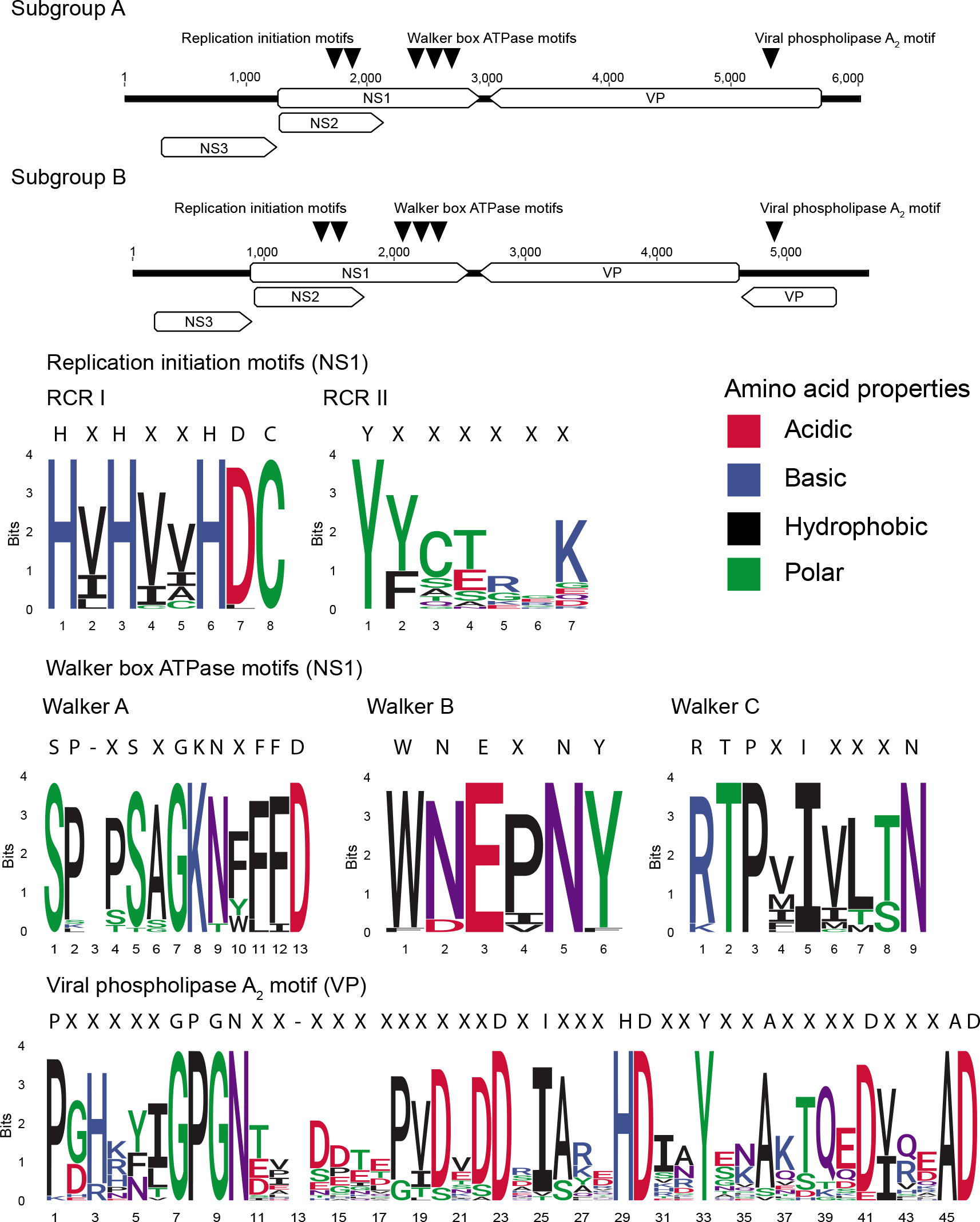
Sea star-associated densoviruses exhibit two genome organizations and contain motifs typical of densoviruses. Triangles indicate position of amino acid motifs, replication initiation motifs, and Walker box ATPase motifs in densoviruses. Consensus sequences above sequence logos are defined by a 90% identity agreement among all sea star-associated densovirus.

### Identification of EVE of densovirus origin

A total of 8 of the 179 transcriptomic libraries contained contigs with densovirus-like sequences. Ten densovirus-like sequences were found among the 8 libraries based homology searches against the sea star-associated densovirus database. Endogenized densovirus elements primarily contained Walker box ATPase motifs with homology to parvoviruses and densoviruses from a broad diversity of host (**Figure 5, Supplemental Table 5**). The transcriptomes containing densovirus-like sequences came from the following species: *Acanthaster planci* (SRA run ID: DRR072325), *Patiria pectinifera* (SRR5229427), *Echinaster spinulosus* (SRR1139455 and SRR2844624), *Acanthaster brevispinus* (SRR276461), *Linckia laevigata* (SRR5438553), and *Asterias rubens* (SRR1139190 and SRR3087891) (**Supplemental Table 4**).

**Figure 5:**
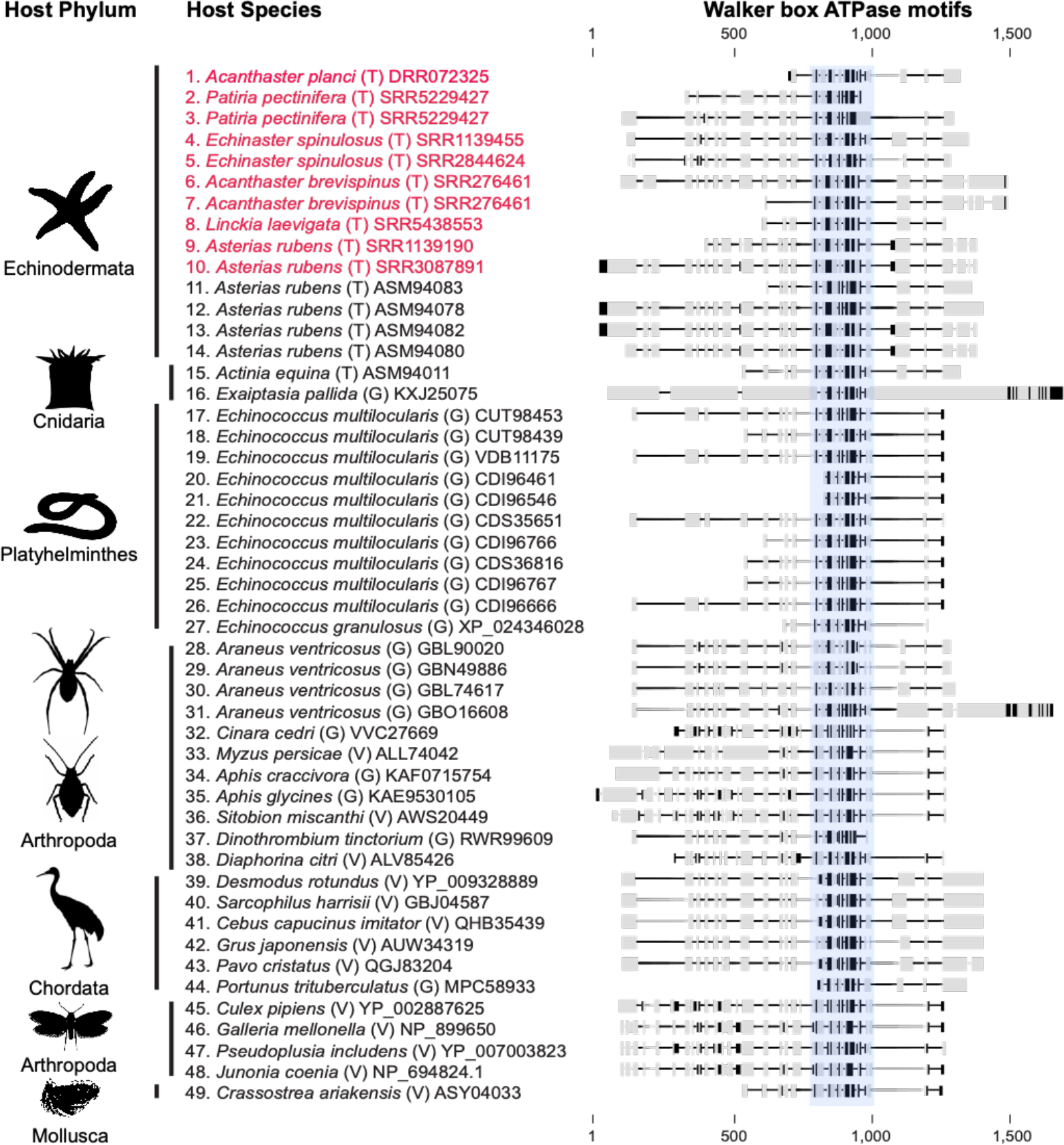
An overview of endogenized densovirus elements (EVE) illustrating the conserved existence of Walker box ATPase motifs. EVEs found in this study are shown in red. Sequences in alignment are grouped by host phylum. Sequences labeled by host species and by origin of sequence from host genome (G), extant virus (V), and host transcriptome (T), and NCBI accession number. Amino acids in bold indicate a sequence identity 75% or greater within the alignment. The blue highlighted region denotes the NTP-binding and helicase region containing Walker A, B, and C motifs found within the NS1/Rep ORF in densoviruses and parvoviruses.

## Discussion Section

The initial investigation for a viral agent associated with SSWS performed viral metagenomic surveys (DNA and RNA) to compare the viral consortia between species and between asymptomatic and symptomatic individuals to determine the most likely viral candidate for further investigation (11). The conclusion from these molecular surveys was that a densovirus, SSaDV, was the sole DNA virus associated with SSWS and was more represented in metagenomic libraries of symptomatic individuals (11). A reanalysis of these data show that SSaDV is one of ten densoviruses present, and shows that SSaDV is neither more abundant by number of reads per library comparing asymptomatic to symptomatic individuals (based on read mapping analysis) nor more prevalent between libraries comparing symptomatic to asymptomatic individuals (**Figure 1, Supplemental Table 3**). This result contradicts the original conclusion that SSaDV is associated with SSWS. SSaDV was, on average, the most abundant densovirus by read mapping analysis and, thus, the most consistently assembled, likely biasing its discovery and the conclusions of previous studies. The higher representation of SSaDV among metagenomic libraries may be the result of higher viral enrichment in those samples prior to sequencing rather than a result of greater viral loads prior to metagenomic preparation. While a higher abundance of SSaDV could reflect important host-virus biology, it remains to be determined whether greater viral loads, determined by qPCR, of SSaDV has any biological significance. A pitfall of viral metagenomics from animal tissue using a viral enrichment method is the significant variability in non-viral genetic material between viromes within a study making quantitative comparisons difficult (33). According to our metagenomic survey, SSaDV is one species within a diverse extant population of densovirus present in sea star populations on the west coast of the United States.

The discovery of densoviruses from sea stars collected from China, Antarctica, and the Pacific and Atlantic coasts of the United States indicates their ubiquitous distribution and substantial extant diversity (**Table 1, Figure 3**). The diversity observed in this study is likely a small fraction of the total diversity among echinoderms, considering these viruses have also been found in sea urchins (14). Sea star associated densoviruses seems to also be pervasive in wild populations. The two densovirus genotypes with the best-described ecological characteristics, SSaDV and AfaDV, share striking similarities. Both viral genotypes are not species-specific, found across a large geographic range, are commonly found in asymptomatic individuals, and have a wide tissue tropism with pyloric caeca being the primary tissue of detection (**Figure 2**) (13). This set of characteristics suggests that both viruses form persistent infections in sea stars.

The genus *Ambidensovirus*, to which both previously described sea star associated densoviruses belonged to based on genome organization, was recently divided into seven newly proposed genera to resolve paraphyly within the genus (8). In this new arrangement, SSaDV and CqDV (*Cherax quadricarinatus* (shrimp) densovirus, the most genetically similar densovirus to SSaDV prior to the discovery of AfaDV) were assigned to the genus *Aquambidensovirus*, putatively uniting all aquatic densoviruses (7, 12, 31). Our phylogenetic analysis did not support the monophyly of sea star associated densoviruses within the newly proposed *Aquambidensovirus* genus, nor did all aquatic densoviruses cluster into a single well-supported clade (**Figure 3**). Newly proposed classification schemes within the *Ambidensovirus* genus would greatly benefit from the inclusion of broader taxonomic sampling before proposing new systematic arrangements of this highly diverse genus.

Given the lack of immortal cell-cultures, the discovery of echinoderm densoviruses has been primarily through metagenomics, and that constraint was the motivation for our analysis of transcriptomes as an additional and alternative option for densovirus discovery. Host transcriptomes have been a rich source of viral discovery from eukaryotes and have expanded our knowledge of host associations for many viral groups (35, 36). However, we did not find transcriptomes to be an effective method for the purpose of densovirus discovery compared to viral metagenomics particularly RNA viral metagenomes. This could be due to various methodological reasons. First, the viral metagenomes prepared for this study were enriched for encapsulated nucleic acids with a cDNA enrichment step; by contrast, transcriptomes target mRNA through rRNA depletion and/or through selection for poly-A tails. Second, to detect DNA viruses from a host transcriptome requires tissue containing an active infection, which may not be detectable without very high sequence depth. The transcribed EVEs found in this study were only detected in transcriptomes larger than 2.4 Gbases. Third, the genomes discovered in the RNA viral metagenomes are likely ssDNA that was carried through the RNA extraction process. ssDNA is an uncommon nucleic acid template for non-viral material and a difficult template to remove during RNA extraction. Most commercial kits use DNases that preferentially target dsDNA and inefficiently cleave ssDNA. Without preferentially targeting mRNA prior to cDNA synthesis in addition to enriching for encapsulated nucleic acid, the chances of picking up ssDNA in a pool of RNA is much higher.

None of the transcriptome-derived densovirus-like sequences appeared to be extant densoviruses based on ORF architecture and motif repertoire. These contigs only encoded part of NS1, lacked RCR motifs, and were not the typical coding length found in sea star-associated densovirus genomes. We conclude that these densovirus-like sequences are likely transcribed EVEs present in host cells. EVEs from *Asterias rubens* and *Acanthaster planci* could be traced to their genomes, while our inability to trace others reflects the lack of publicly available host genomes. The putative EVEs present in *Asterias rubens* were nearly identical to those previously reported in the same host (37) though most we observed had low sequence identity to previously identified EVEs from other invertebrates. It is likely that these EVEs have been established in the germline of *Asterias rubens*, and our findings corroborate previous work proposing sea star densoviruses can infect germ line cells (13). The expression of these EVEs in *Asterias rubens* was found to trigger the RNA interference (RNAi) response, specifically the Piwi-dependent pathway, signifying these EVEs are still recognized as foreign and are regulated through the immune system (37). This RNAi response has been widely observed in terrestrial invertebrate genomes containing EVEs descending from densovirus (38). We expect that the expansion of echinoderm genomes, and corresponding small RNA libraries, will further support this conclusion.

Essentially all observed EVEs retained the Walker Box ATPase motifs which collectively function as a helicase (**Figure 5**) (39). This helicase domain belongs to the superfamily III helicases (SF3) which are more broadly grouped as AAA + ATPases (40). SF3 helicases are only encoded by DNA and RNA viruses so their presence in cellular genomes must be the result of endogenization (41, 42). The retention of the Walker box ATPase motifs among endogenized densovirus elements has been observed across a diverse range of invertebrate hosts, suggesting a beneficial function for coopting and possibly maintaining the function of the SF3 helicase (10, 12, 38, 43, 44). The adaptive benefit of a Walker Box ATPase-containing EVE has been demonstrated in the pea aphid (*Acyrthosiphon pisum*), where wing development was regulated by two modified densovirus NS1 EVEs, which only retained the Walker Box ATPase motifs (44). The expression of these two EVEs in crowded conditions initiated wing development, which could be suppressed by knocking down their expression. These results demonstrate that this viral gene can be co-opted by the host to modulate the response of a phenotypically plastic trait to environmental cues. Another plausible hypothesis for EVE function is the ability to enhance or prime the immune system against new infections (38). However, we observed little sequence identity between extant sea star-associated densoviruses and the EVEs in their transcriptomes. This sequence differences suggest that these EVES are unlikely to have a role in priming the piRNA response against new infections.

We employed a metagenomic and transcriptomic approach to explore the diversity of sea stars associated-densoviruses, while advancing understanding of the biogeography of SSaDV, the first densovirus found in sea stars. Empirically, we found that viral metagenomes provided a more effective resource for densovirus discovery compared to host transcriptomes. We discovered 37 new densovirus genomes from sea stars and identified EVEs expressed in host transcriptomes that are of densovirus origin based on detection of the tripartite SF3 helicase domain in these EVEs. Using PCR, we found SSaDV to have a putatively wide tissue tropism, with the pyloric caeca being the most consistent tissue for viral detection. SSaDV was detected across a broad latitudinal range in the northeastern Pacific from southern California to Alaska and found in tissues in nearly all sea star species tested. These results corroborate the hypothesis that these viruses are common among populations and suggest they form persistent infections in sea stars. Given the diversity of densoviruses and their broad distribution among tissues, populations, and species of both healthy and diseased sea stars, we propose that the association of SSaDV with Sea Star Wasting Syndrome should be critically reassessed relative to the mounting evidence that this virus may not be pathogen that causes this disease and instead a common constituent of these animals’ microbiomes.

## Acknowledgments

The authors thank Dr. Sarah Gravem (Oregon State University), Dr. Bruce Menge (Oregon State University), Dr. Lauren Schiebelhut (UC Davis), Dr. Michael Dawson (UC Merced), Elizabeth Ashley (SeaDoc Society), Erika Nilson (SeaDoc Society), Dr. Melissa Pespeni (University of Vermont), Dr. John Ware (University of Georgia), Christoph Pierre (UC Santa Barbara), Carter Urnes (National Park Service), Dr. Steven Fradkin (National Park Service), Dr. Nathalie Oulhen (Brown University), Dr. Gary Wessel (Brown University), Dr. Sarah Cohen (San Francisco State University), Dr. Margaret Amsler (University of Alabama), Dr. James McClintock (University of Alabama), Dr. Chuck Amsler University of Alabama), and Sean Williams (Hoonah Indian Association) for assistance in sample collection.

## Table Legends

Table 1: Sea star-associated densovirus genome characteristics and metadata

Supplemental Table 1: Metadata for tissue samples collected for DNA extraction and PCR testing for SSaDV

Supplemental Table 2: Viral metagenomes analyzed for densovirus sequences.

Supplemental Table 3: Read mapping analysis of newly discovered densovirus genomes from viral metagenomes prepared in Hewson et al., 2014.

Supplemental Table 4: Sea star transcriptomes surveyed for densovirus sequences

Supplemental Table 5: Top BLASTp result against the NCBI non-redundant database querying putative densovirus-like elements

