## Supplemental Table 3 for "Diversity of sea star-associated densoviruses and transcribed endogenized viral elements of densovirus origin"

| Library | Host | Collection Site | Metagenome | Health | PhaDV1 | PhaDV2 | PhaDV3 | PhaDV4 | PhaDV5 | PhaDV6 | EtaDV1 | EtaDV2 | EtaDV3 | SSaDV | Total Reads | Library Reads | Percent Reads |
| --- | --- | --- | --- | --- | --- | --- | --- | --- | --- | --- | --- | --- | --- | --- | --- | --- | --- |
| V01 | *Dermasterias imbricata* | Friday Harbor, WA | DNA | Diseased | 0 | 0 | 0 | 5 | 0 | 0 | 0 | 0 | 0 | 1 | 6 | 1003976 | 5.98E-06 |
| V02 | *Solaster stimpsoni* | Friday Harbor, WA | DNA | Diseased | 0 | 0 | 0 | 0 | 0 | 0 | 0 | 0 | 0 | 3 | 3 | 3487142 | 8.60E-07 |
| V03 | *Solaster stimpsoni* | Friday Harbor, WA | DNA | Diseased | 0 | 0 | 0 | 0 | 0 | 0 | 0 | 0 | 0 | 2 | 2 | 1450512 | 1.38E-06 |
| V04 | *Evasterias troschelii* | Friday Harbor, WA | DNA | Diseased | 0 | 0 | 0 | 0 | 0 | 0 | 0 | 0 | 0 | 0 | 0 | 1943056 | 0 |
| V05 | *Pycnopodia helianthoides* | Friday Harbor, WA | DNA | Diseased | 0 | 0 | 0 | 0 | 0 | 0 | 0 | 0 | 0 | 0 | 0 | 3221364 | 0 |
| V06 | *Evasterias troschelii* | Friday Harbor, WA | DNA | Healthy | 0 | 0 | 0 | 0 | 0 | 0 | 0 | 0 | 0 | 1 | 1 | 1427450 | 7.01E-07 |
| V07 | *Pycnopodia helianthoides* | Friday Harbor, WA | DNA | Healthy | 0 | 0 | 0 | 0 | 0 | 0 | 0 | 0 | 0 | 0 | 0 | 676082 | 0 |
| V08 | *Pycnopodia helianthoides* | Burrard Inlet BC | DNA | Healthy | 6 | 7 | 0 | 6 | 0 | 0 | 33 | 0 | 0 | 10334 | 10386 | 1915648 | 0.005421664 |
| V09 | *Pisaster ochraceus* | Burrard Inlet, BC | DNA | Healthy | 0 | 0 | 0 | 0 | 0 | 0 | 0 | 0 | 0 | 14 | 14 | 1176524 | 1.19E-05 |
| V10 | *Pycnopodia helianthoides* | Seattle Aquarium, WA | DNA | Healthy | 3 | 0 | 0 | 0 | 0 | 0 | 0 | 0 | 0 | 2 | 5 | 2761526 | 1.81E-06 |
| V11 | *Pycnopodia helianthoides* | Seattle Aquarium, WA | DNA | Healthy | 0 | 0 | 3 | 4 | 0 | 0 | 0 | 0 | 0 | 2 | 9 | 1460518 | 6.16E-06 |
| V12 | *Pycnopodia helianthoides* | Seattle Aquarium, WA | DNA | Diseased | 27 | 0 | 0 | 21 | 0 | 14 | 0 | 0 | 0 | 13 | 75 | 3267472 | 2.30E-05 |
| V13 | *Pycnopodia helianthoides* | Seattle Aquarium, WA | DNA | Diseased | 0 | 8 | 3 | 17 | 2 | 3 | 0 | 0 | 2 | 5 | 40 | 1617558 | 2.47E-05 |
| V14 | *Pisaster ochraceus* | Olympic National Park, WA | DNA | Healthy | 0 | 0 | 0 | 0 | 0 | 2 | 0 | 0 | 0 | 0 | 2 | 1255710 | 1.59E-06 |
| V15 | *Pisaster ochraceus* | Olympic National Park, WA | DNA | Healthy | 0 | 0 | 0 | 0 | 0 | 0 | 0 | 0 | 0 | 0 | 0 | 1225146 | 0 |
| V16 | *Pisaster ochraceus* | Olympic National Park, WA | DNA | Diseased | 0 | 2 | 0 | 0 | 0 | 0 | 0 | 0 | 0 | 0 | 2 | 794834 | 2.52E-06 |
| V17 | *Pisaster ochraceus* | Olympic National Park, WA | DNA | Diseased | 0 | 0 | 0 | 0 | 0 | 0 | 0 | 0 | 0 | 0 | 0 | 1246682 | 0 |
| V18 | *Pisaster ochraceus* | Santa Cruz, CA | DNA | Diseased | 14 | 0 | 0 | 0 | 0 | 6 | 0 | 0 | 0 | 0 | 20 | 1227630 | 1.63E-05 |
| V19 | *Pisaster ochraceus* | Santa Cruz, CA | DNA | Diseased | 40 | 0 | 0 | 3 | 0 | 2 | 0 | 0 | 0 | 0 | 45 | 1497116 | 3.01E-05 |
| V20 | *Pisaster ochraceus* | Santa Cruz, CA | DNA | Healthy | 73 | 0 | 0 | 0 | 0 | 3 | 0 | 0 | 0 | 0 | 76 | 1919002 | 3.96E-05 |
| V21 | *Pisaster ochraceus* | Santa Cruz, CA | DNA | Healthy | 0 | 0 | 0 | 1 | 0 | 0 | 0 | 0 | 0 | 0 | 1 | 1130796 | 8.84E-07 |
| V22 | *Pycnopodia helianthoides* | Gower Point, BC | DNA | Diseased | 0 | 0 | 0 | 0 | 0 | 0 | 0 | 0 | 2 | 2 | 4 | 1399640 | 2.86E-06 |
| V23 | *Pycnopodia helianthoides* | Gower Point, BC | DNA | Healthy | 0 | 0 | 0 | 0 | 0 | 0 | 2 | 0 | 3 | 8 | 13 | 1798996 | 7.23E-06 |
| V24 | *Evasterias troschelii* | Cape Roger Curtis, BC | DNA | Diseased | 0 | 0 | 0 | 3 | 0 | 0 | 176 | 0 | 38 | 128 | 345 | 2602550 | 0.000132562 |
| V25 | *Evasterias troschelii* | Cape Roger Curtis, BC | DNA | Diseased | 0 | 4 | 0 | 2 | 0 | 0 | 107 | 2 | 21 | 45 | 181 | 1064000 | 0.000170113 |
| V26 | *Evasterias troschelii* | Cape Roger Curtis, BC | DNA | Healthy | 4 | 32 | 6 | 76 | 0 | 7 | 314 | 1550 | 748 | 848 | 3585 | 642152 | 0.00558279 |
| V27 | *Evasterias troschelii* | Cape Roger Curtis, BC | DNA | Healthy | 0 | 0 | 0 | 0 | 0 | 0 | 2 | 8 | 7 | 22 | 39 | 459414 | 8.49E-05 |
| V28 | *Pitiria miniata* | Vancouver Aquarium,BC | DNA | Diseased | 10 | 0 | 0 | 0 | 0 | 0 | 0 | 0 | 0 | 0 | 10 | 1043014 | 9.59E-06 |
| V30 | *Pycnopodia helianthoides* | Seattle Aquarium, WA | RNA | Healthy | 0 | 0 | 0 | 1 | 0 | 0 | 18 | 0 | 0 | 1 | 20 | 4190866 | 4.77E-06 |
| V31 | *Pycnopodia helianthoides* | Seattle Aquarium, WA | RNA | Healthy | 653 | 22 | 219 | 658 | 50 | 66 | 47 | 32 | 58 | 316 | 2121 | 4049538 | 0.000523763 |
| V32 | *Pycnopodia helianthoides* | Seattle Aquarium, WA | RNA | Diseased | 26 | 11 | 0 | 29 | 0 | 15 | 2 | 0 | 0 | 48 | 131 | 1519942 | 8.62E-05 |
| V33 | *Pycnopodia helianthoides* | Seattle Aquarium, WA | RNA | Diseased | 710 | 685 | 106 | 1027 | 213 | 385 | 4 | 0 | 14 | 601 | 3745 | 2377178 | 0.001575397 |
