## Supplemental Table 5 for "Diversity of sea star-associated densoviruses and transcribed endogenized viral elements of densovirus origin"

| Contig ID | Contig Length (nucleotide) | Parvovirus ORF Length (amino acid) | Complete or Partial ORF | Adjacent transposable element | Found in Host Genome | Query Library | Sea star species | BLASTp ID | GenBank Accession | e-value | % ID | Query Cover |
| --- | --- | --- | --- | --- | --- | --- | --- | --- | --- | --- | --- | --- |
| DN57314 | 938 | 312 | Partial | No | Yes | DRR072325 | Acanthaster planci | hypothetical protein AVEN_50382_1 [Araneus ventricosus] | GBO16608.1 | 8.00E-32 | 40% | 54% |
| DN11764 | 3,031 | 593 | Complete | No | Yes | SRR1139190 | Asterias rubens | putative non-structural NS1 [Caledonia starfish parvo-like virus 1] | ASM94080.1 | 0.00E+00 | 99.83% | 100% |
| DN11764 | 3,031 | 311 | Partial | No | Yes | SRR1139190 | Asterias rubens | putative structural VP [Caledonia starfish parvo-like virus 1] | ASM94081.1 | 0.00E+00 | 100% | 100% |
| DN8030 | 2,908 | 614 | Complete | No | No Genome Available | SRR1139455 | Echinaster spinulosus | NS1-like protein [Dinothrombium tinctorium] | RWR99609.1 | 2.00E-46 | 32.77% | 57% |
| DN6488 | 1,501 | 466 | Complete | No | No Genome Available | SRR2844624 | Echinaster spinulosus | NS1-like protein [Dinothrombium tinctorium] | RWR99609.1 | 2.00E-47 | 32.68% | 75% |
| DN27874 | 2,686 | 827 | Complete | No | Yes | SRR3087891 | Asterias rubens | putative non-structural NS1 [Caledonia starfish parvo-like virus 2] | ASM94082.1 | 0.00E+00 | 90.33% | 100% |
| DN6783 | 2,299 | 544 | Partial | No | No Genome Available | SRR5229427 | Patiria pectinifera | hypothetical protein AVEN_50382_1 [Araneus ventricosus] | GBO16608.1 | 3.00E-26 | 27.15% | 65% |
| DN46266 | 1,013 | 328 | Partial | No | No Genome Available | SRR5229427 | Patiria pectinifera | NS1-like protein [Dinothrombium tinctorium] | RWR99609.1 | 9.00E-26 | 36.55% | 57% |
| DN158628 | 1055 | 340 | Complete | No | No Genome Available | SRR5438553 | Linckia laevigata | putative non-structural NS1 [Caledonia starfish parvo-like virus 3] | ASM94083.1 | 7.00E-81 | 38.44% | 96% |
| DN57645 | 3,411 | 880 | Complete | No | No Genome Available | SRR9276461 | Acanthaster brevispinus | putative non-structural NS1 [Caledonia starfish parvo-like virus 1] | ASM94080.1 | 5.00E-131 | 35.52% | 76% |
| DN63527 | 16,517 | 462 | Complete | Yes | No Genome Available | SRR9276461 | Acanthaster brevispinus | putative non-structural NS1 [Caledonia starfish parvo-like virus 3] | ASM94083.1 | 1.00E-61 | 35.20% | 69% |
